# Movement planning predictions shape somatosensory sensitivity

**DOI:** 10.64898/2026.06.12.731893

**Authors:** Pierangelo Nicolas D’Onofrio Pacheco, Eckart Zimmermann

## Abstract

Tactile sensitivity is reduced during limb movement, a phenomenon known as somatosensory gating. Here, across two experiments, we ask whether gating is driven by motor prediction or by motor execution. In a Go/NoGo paradigm, in which participants planned movements in every trial but on rare trials had to withhold them. Since previous work indicated that predictions about movement kinematics influence gating also during passive movements, we also tested a mechanical arm transport in a passive Go/NoGo paradigm. Perceived intensity was attenuated exclusively on Go trials, in both active and passive movements, and was indistinguishable from baseline on NoGo trials. However, discrimination precision was selectively degraded in Active NoGo trials when a movement was planned and then withheld. In Experiment 2, vibro-tactile probes delivered before movement onset already showed the same bias as probes delivered during movement, in both active and passive conditions, while precision remained unchanged. Our data demonstrates that the sensorimotor system predictively establishes tactile precision before movement onset. Such a mechanism might contribute to active texture exploration by separating tactile signals from the sensory signals arising through the self-produced movement.

## Introduction

Tactile stimuli delivered during limb movements are often perceived as less intense than stimuli presented at rest, a phenomenon known as somatosensory gating. Since its initial description, there has been considerable debate as to whether this reduction reflects peripheral masking and other bottom-up factors or an active modulation of sensory processing associated with movement generation(1).

We have recently found that felt tactile stimulation intensity and discrimination sensitivity are differentially affected during arm movements (2) (3). Consistent with other reports, we found that the felt intensity is reduced likewise during active and passive movements(4–7), suggesting that attenuation of stimulus intensity does not depend on motor planning. In contrast, tactile discrimination sensitivity depended strongly on movement predictability. Discrimination performance was impaired during passive movements when movement speed varied unpredictably across trials, whereas sensitivity remained high during active movements and during passive movements with constant, predictable speed. These findings suggest that tactile sensitivity is not generally suppressed during movement but can be preserved when the sensory consequences of movement are predictable.

The reduction in felt intensity during movement has also been linked to the anticipation of grasping control (8). Somatosensory gating increases when movement-relevant object properties, such as mass distribution, are predictable, suggesting that sensory attenuation depends on how well object features can be anticipated. Somatosensory gating selectively reduced the vibration frequency that matched the vibrotactile feedback when participants stroked across textured objects (9). More generally, studies of tactile detection and suppression during reach-to-grasp tasks confirm that gating depends on task relevance and predicted sensory consequences(10, 11)

The link between tactile precision and movement prediction may be particularly relevant for tactile texture perception. Humans typically assess surface properties through active exploratory movements, during which the tactile signals generated at the skin depend strongly on movement parameters such as scanning speed and contact force (12). Consequently, the nervous system must distinguish between sensory signals arising from the exploratory movement itself and those that convey information about surface properties. Despite the movement dependence of tactile texture signals, texture perception remains remarkably robust across variations in exploratory movements, including changes in scanning speed (13). This perceptual invariance suggests the existence of mechanisms that compensate for predictable movement-related sensory input while preserving task-relevant tactile information.

Here, we aimed to study separately both the influence of movement planning and the influence of movement execution. To directly demonstrate how movement planning influences tactile perception, it is necessary to separate planning processes from movement execution. A useful experimental approach for achieving this dissociation is the go–no-go paradigm. In this setup, participants plan to perform a movement, but on a small number of trials they receive a late cue instructing them to withhold the movement just before execution.

If somatosensory gating reflects an influence of motor planning of sensory processing, then it should be engaged whenever the motor system is set up for an upcoming action, regardless of whether that action is ultimately executed. If instead it reflects a consequence of the motor act itself or reafferent input, then foreknowledge alone should not suffice. The separate influence of movement planning and execution on tactile perception has been examined by several studies (14, 15). Voss et al. (2008) (16) delivered electrical stimuli 50 ms before the appearance of a cue which instructed which finger to move. They found that under asymmetric cue probabilities (80:20), stimuli were attenuated even on trials where the non-expected finger was ultimately cued. Voudouris and Fiehler (2017) (17) measured both bias and precision during reaching and found that changes appeared only during movement execution, not during the planning phase.

Another question concerns the time-course of somatosensory gating. Somatosensory-evoked potentials are attenuated during the period preceding voluntary action (18) (19) (20) (21), and vibro-tactile detection can be gated before movement in bimanual reaching tasks (22). Using TMS to delay the cortical output of motor commands, Voss et al. (2006) (23) showed that somatosensory gating during the post-TMS silent period was comparable to that during actual movement, demonstrating that central motor signals upstream of primary motor cortex are sufficient for attenuation even in the temporary absence of overt motion. However, most studies of pre-movement somatosensory gating have quantified either detection thresholds, detection rates, or evoked responses without separating bias and discriminability within a single intensity discrimination task. It therefore remains unclear whether pre-movement somatosensory gating is broad and degrading tactile processing in general or selective, primarily shifting perceived intensity while leaving discrimination intact.

Here we ask whether somatosensory gating is driven by movement execution or motor planning. Using a Go/NoGo paradigm (Experiment 1) and a pre-movement timing paradigm (Experiment 2), we tested whether perceived intensity and discrimination precision are differently sensitive to motor execution, motor planning, and violatons of motor predictions.

## Methods

### Apparatus

Participants sat in front of a 24-inch monitor (1920 × 1080 px, 144 Hz refresh rate) at 57 cm. The right arm rested on an ergonomic support. In the active conditions, participants moved a Touch X haptic stylus (3D Systems) along a 20 cm horizontal path between two visual targets displayed on the screen. In the passive conditions, the stylus was attached to a custom linear rail driven by a stepper motor (Arduino Leonardo controller) that translated the hand at a constant velocity of 200 mm/s.

Tactile stimulation was delivered by a 1 cm vibro-tactile actuator fixed over the median-nerve region of the right forearm. Each vibration lasted 300 ms. On every trial, the reference vibration had a fixed duty cycle of 55%, while the comparison vibration was randomly selected from nine possible duty cycles (10–90% in 10% increments). Vibrations were triggered by an Arduino Nano microcontroller synchronized with the experimental timing routines. Participants wore Soundcore Life Q30 noise-canceling headphones to mask auditory cues from the movement apparatus. Responses were given on a three-key keypad (Pimoroni Keybow) by pressing “1” if the first vibration felt stronger or “2” if the second did. All experimental measurements including stimulus presentation, stylus tracking, stimulus timing, and response collection were controlled by custom C# scripts running in Unity.

Responses were fitted with cumulative Gaussian psychometric functions using maximum-likelihood estimation for each participant and condition. The point of subjective equality (PSE) indexed perceptual bias, with lower values indicating reduced perceived intensity. The just-noticeable difference (JND) was given by the standard deviation of the fitted function, indexing discrimination precision, with lower values indicating higher sensitivity. Trials with missing or unusable responses were excluded prior to fitting. Participants with more than 25% missing data were excluded from analysis. All statistical analyses were performed in JASP 0.18.1.0, with α = .05. Post-hoc comparisons used Holm-Bonferroni correction. Effect sizes are reported as partial η² for ANOVAs and Cohen’s d for t-tests.

### Experiment 1: Go/NoGo gating

#### Participants

21 right-handed adults were recruited. The study conformed to the Declaration of Helsinki and was approved by the ethics committee of the Faculty of Mathematics and Natural Sciences, Heinrich-Heine University Düsseldorf. All participants provided written informed consent prior to participation. Two participants were excluded due to excessive missing data, resulting in a final sample of 19 (mean age = 25.31 years, SD = 3.4) participants.

#### Design and procedure

The experiment followed a 2 (Movement type: Active, Passive) × 2 (Trial type: Go, NoGo) factorial within-subjects design (Figure 2), with an additional stationary control condition. Active and passive trials were presented blockwise, with block order randomized across participants.

Each trial began when the participant positioned the stylus inside the start sphere on the left (see Figure 1 or 2), a red sphere of 2 cm diameter displayed on the screen which turned yellow on touch indicating the trial onset. After 1000 ms, the target sphere, a second sphere of identical size, positioned to the right such that the linear path from the center of the start sphere to the center of the target sphere measured 20 cm turned green, signaling the participant to execute the movement. On Go trials (70% of trials), the target sphere remained green and the participant executed the movement. On NoGo trials (30% of trials), the target sphere turned red 100 ms after the green onset, instructing the participant to withhold the movement and remain stationary. Trial type was randomized within each block.

**Figure 1.**
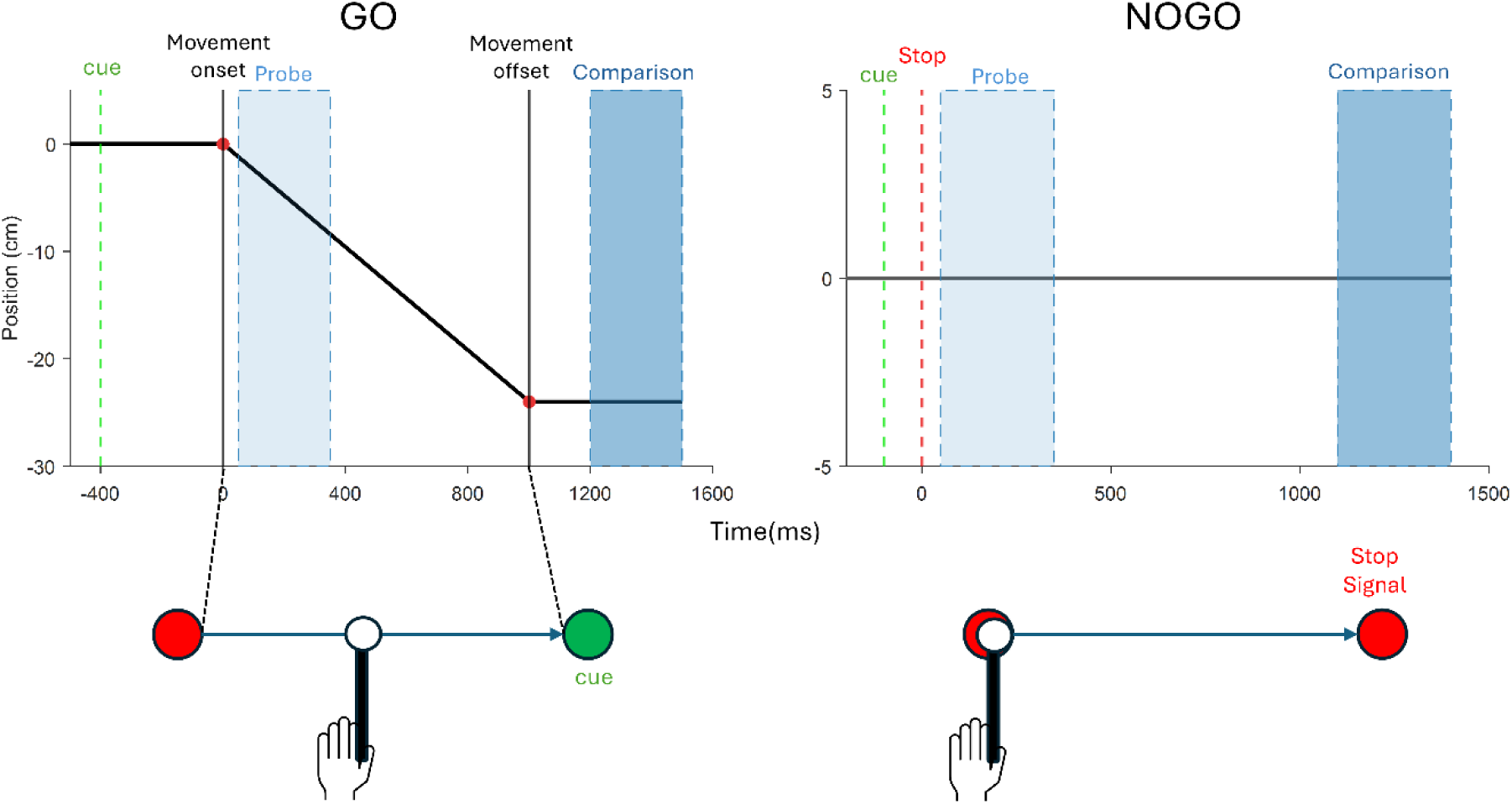
**Experimental paradigm**: Experiment 1 (Go/NoGo). Trial timeline for Go (left) and NoGo (right) conditions. The upper panels show arm position (cm) as a function of time (ms). On Go trials, the green cue triggered movement execution; the reference vibration (Probe) was delivered 200 ms after movement onset, and the comparison vibration was delivered 500 ms after reaching the target. On NoGo trials, a red stop signal appeared 100 ms after the green cue; the probe was delivered 50 ms after the stop signal while the arm remained stationary, and the comparison followed 800 ms later. The lower panels illustrate the corresponding screen display: the red start sphere, the white intermediate position, and the green (Go) or red (NoGo) target sphere, with the stylus held by the participant.

**Figure 2.**
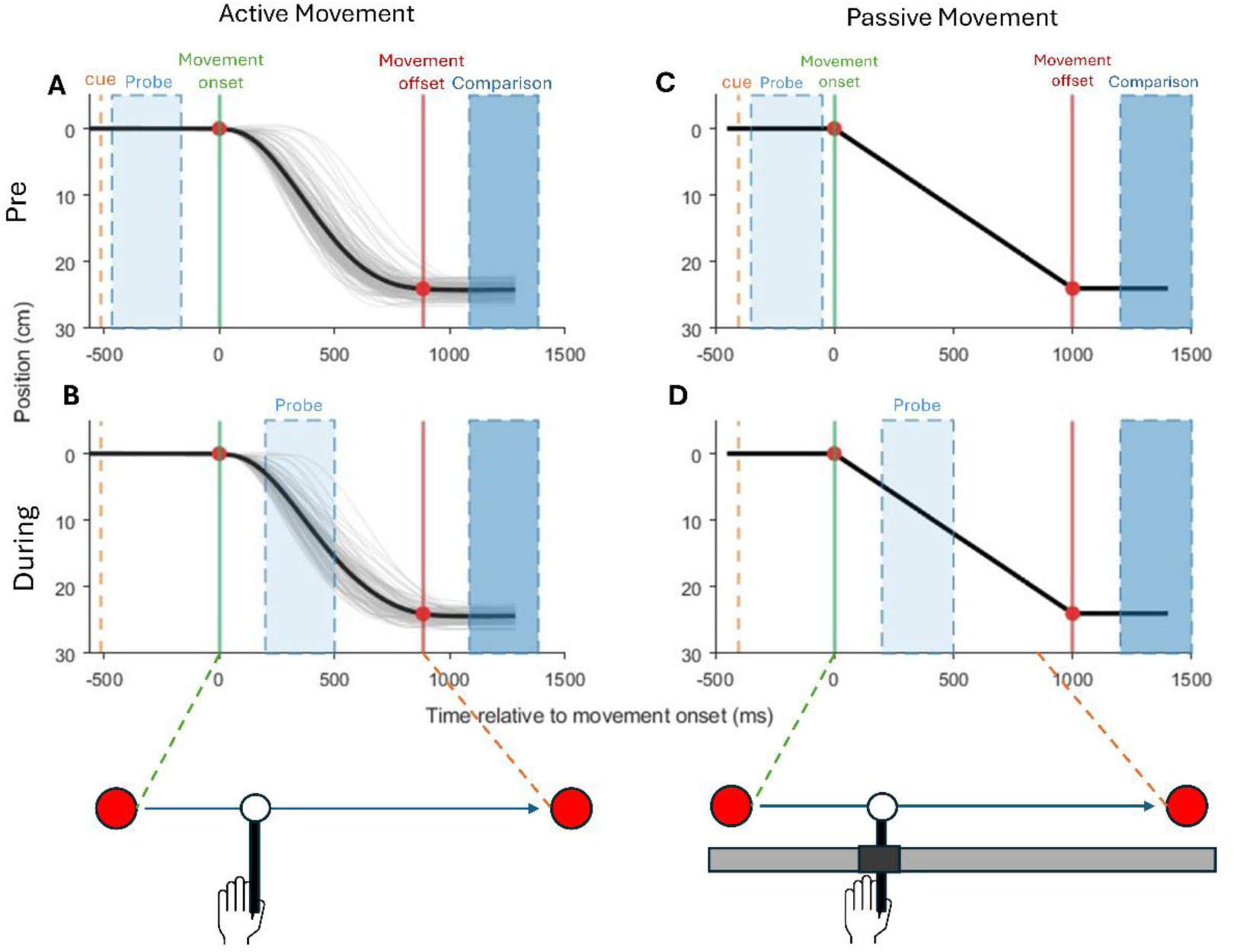
**Experimental paradigm**: Experiment 2 (Pre-movement). Trial timelines for Active (left panels, A–B) and Passive (right panels, C–D) conditions, shown separately for pre-movement (top row, A and C) and During-movement (bottom row, B and D) probe delivery. Position (cm) is plotted as a function of time relative to movement onset (ms). Grey lines show individual participant trajectories; the black line shows the group mean. The orange dashed line marks the visual cue; the green line marks movement onset; the red line marks movement offset. The light blue shaded region indicates the probe window; the dark blue shaded region indicates the comparison window. The lower diagrams illustrate the screen display for active (left, hand-held stylus) and passive (right, motorized rail) conditions.

*Go trials.* In the active Go condition, participants moved the stylus from the start sphere to the target sphere along the 20 cm horizontal path. The reference vibration was delivered 200 ms after the participant exited the start sphere, during movement execution. In the passive Go condition, the motorized rail-initiated arm displacement 200 ms after the green cue, and the reference vibration was likewise delivered 200 ms after leaving the start position during passive displacement. In both conditions, the comparison vibration was delivered 500 ms after reaching the target sphere, see figure 1 left panel.

*NoGo trials.* In both active and passive NoGo conditions, the target sphere first turned green as in Go trials, but turned red 100 ms after the green onset, instructing the participant to withhold movement. The reference vibration was delivered with a 50 ms delay after the red stop signal. Because the vibration lasted 300 ms and movement had been inhibited, stimulation occurred entirely while the arm was stationary. The comparison vibration was delivered 800 ms after the reference vibration. Participants made the same intensity judgment while remaining in the start position. If a participant exited the start sphere during a NoGo trial, the trial was flagged as invalid and excluded from analysis. See figure 1 on the right panel.

##### Control condition

The arm remained stationary, and participants received the reference followed by the comparison 500 ms later.

Participants completed three blocks of 90 trials for each movement condition and one control block of 90 trials, yielding a total of 630 trials per participant. Within each movement block, approximately 63 trials were Go and 27 were NoGo.

#### Analysis

PSE and JND values were analyzed with one-way repeated-measures ANOVAs across the five conditions (Control, Active Go, Active NoGo, Passive Go, Passive NoGo). To directly quantify the effect of movement cancellation on precision, we computed for each participant the JND difference between Go and NoGo (Delta = Go − NoGo) separately for active and passive conditions. Delta scores were tested against zero with one-sample t-tests and compared between movement types with a paired-samples t-test.

### Experiment 2: Pre-movement gating

#### Participants

Twenty-five right-handed adults were recruited. All reported normal or corrected-to-normal vision and no history of neurological or psychiatric disorders. One participant was excluded due to excessive missing data, resulting in a final sample of twenty-four participants (19 females; mean age = 21.3 years, SD = 2.1). The study conformed to the Declaration of Helsinki and was approved by the ethics committee of the Faculty of Mathematics and Natural Sciences, Heinrich-Heine University Düsseldorf. All participants provided written informed consent prior to participation.

#### Design and procedure

The experiment followed a 2 (Condition: Active, Passive) × 2 (Time: Pre-movement, during movement) factorial within-subjects design (Figure 1), participants also completed an additional stationary control condition. Block order was randomized across participants, with each block containing exclusively active or passive trials.

Each trial began with the appearance of two-colored spheres on the screen corresponding to the start and endpoint of the horizontal movement path. After 1000ms the color of the second sphere changed, serving as the movement cue, signaling participants to initiate the movement (active) or indicating the imminent onset of passive displacement. See figure 2.

During the normal trials (75% of trials per block), the reference vibration was delivered 200 ms after movement onset in both the active and passive conditions. On pre-movement trials (25% of trials per block), the reference vibration was delivered 50 ms after the visual cue, before movement had begun (mean=601ms, SD=193ms). In the active condition, this corresponded to the movement planning phase preceding voluntary movement initiation (figure 2, A & B panels). In the passive condition, the device initiated arm displacement 300 ms after the visual cue, so the pre-movement probe was delivered 250 ms before movement onset (figure 2, C & D panels). Trial type was randomized within each block. In both conditions, the comparison vibration was delivered 200 ms after the arm reached the endpoint. In the control condition, the arm remained stationary throughout; participants received the reference followed by the comparison 500 ms later.

Participants completed four blocks for each movement condition, with 90 trials per block, yielding 360 active trials (90 pre-movement, 270 during movement) and 360 passive trials (90 pre-movement, 270 during movement), plus one control block of 90 trials, for a total of 810 trials.

#### Analysis

PSE and JND values were analyzed with a 2 (Condition: Active, Passive) × 2 (Time: Pre-movement, During movement) repeated-measures ANOVA. Additional one-way repeated-measures ANOVAs compared each movement condition against the stationary control. A five-level one-way ANOVA across all conditions was also performed. Additionally, active pre-movement trials in which movement onset preceded vibration offset (latency < 350 ms) were excluded to ensure that stimulation occurred entirely before movement began (11.2% of pre-movement trials excluded on average).

## Results

### Experiment 1: Go/NoGo gating

#### Bias (PSE)

A one-way repeated-measures ANOVA across the five conditions revealed a significant main effect, F(4, 72) = 16.48, p < .001. Post-hoc comparisons revealed that perceived intensity was attenuated exclusively on Go trials. (figure 3 A)

**Figure 3.**
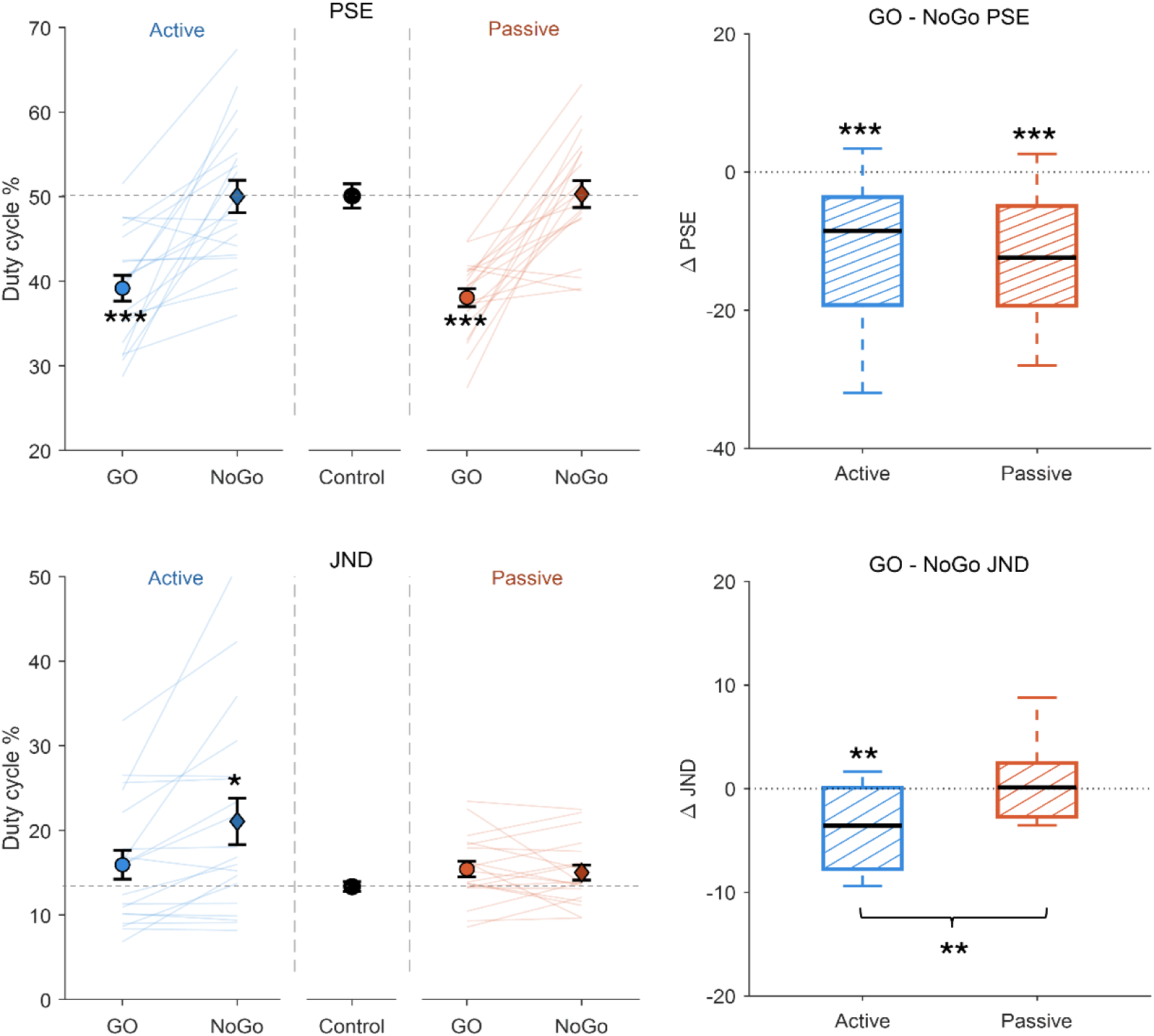
**Results**: Experiment 1 (Go/NoGo). Left panels show individual and mean PSE (top) and JND (bottom) values in duty cycle (%) for Active (blue) and Passive (orange) conditions across Go, NoGo, and Control. Thin lines connect individual participants across conditions. Error bars represent ±1 SEM. The dashed grey horizontal line indicates the reference duty cycle (55% for PSE, mean control JND for JND). Asterisks indicate significant deviation from Control (one-sample tests; *p < .05, **p < .01, ***p < .001). Right panels show control-normalized difference scores (Δ = Go − NoGo) for PSE (top) and JND (bottom). Boxplots show median, interquartile range, and whiskers to 1.5×IQR. The bracket in the lower right panel indicates a significant difference between Active and Passive delta JND. **p < .01, ***p < .001.

Both Go conditions differed significantly from Control: Active Go (mean difference = 1.091, p < .001) and Passive Go (mean difference = 1.419, p < .001). Neither NoGo condition differed from Control: Active NoGo (mean difference = −0.082, p = 1.000) and Passive NoGo (mean difference = −0.118, p = 1.000). The two NoGo conditions did not differ from each other (mean difference = −0.036, p = 1.000), confirming that both were indistinguishable from the stationary baseline. Go and NoGo conditions differed significantly within each movement type: Active Go vs. Active NoGo (mean difference = −1.173, p = .001) and Passive Go vs. Passive NoGo (mean difference = −1.537, p = .002). Active Go and Passive Go did not differ from each other (p = .807). See figure 3B.

#### Precision (JND)

A one-way repeated-measures ANOVA on JND values revealed a significant main effect, F(4, 72) = 4.23, p = .004. The pattern differed markedly from that observed for PSE. Post-hoc comparisons showed that precision was selectively degraded in the Active NoGo condition: Active NoGo JND was significantly higher than Control (p = .049) and significantly higher than Active Go (p = .036). Passive Go and Passive NoGo were virtually identical (mean difference = 0.006, p = 1.000), and neither differed significantly from Control. (figure 3C)

#### Movement cancellation and precision: delta analysis

One sample t-tests on the JND difference scores (Delta = Go − NoGo) confirmed a significant precision cost of cancellation in the active context: Delta (Active) differed significantly from zero, t(18) = −3.303, p = .004. Delta (Passive) did not differ from zero, t(18) = 0.047, p = .963. A paired-samples t-test confirmed that the two deltas differed significantly, t (18) = −3.549, p = .002, establishing a selective precision cost of cancelling a voluntary motor plan that was absent when passive displacement was withheld. See Figure 3D.

### Experiment 2: Pre-movement gating

#### Bias (PSE)

A 2 × 2 repeated-measures ANOVA on PSE values revealed a significant main effect of Condition, F(1, 23) = 17.13, p < .001, η²p = .43, but no main effect of Time, F(1, 23) = 1.11, p = .302, and no Condition × Time interaction, F(1, 23) = 1.19, p = .287.PSE values were lower (greater attenuation) in the passive than in the active context (mean difference = 0.596, p < .001). See Figure 4A.

**Figure 4.**
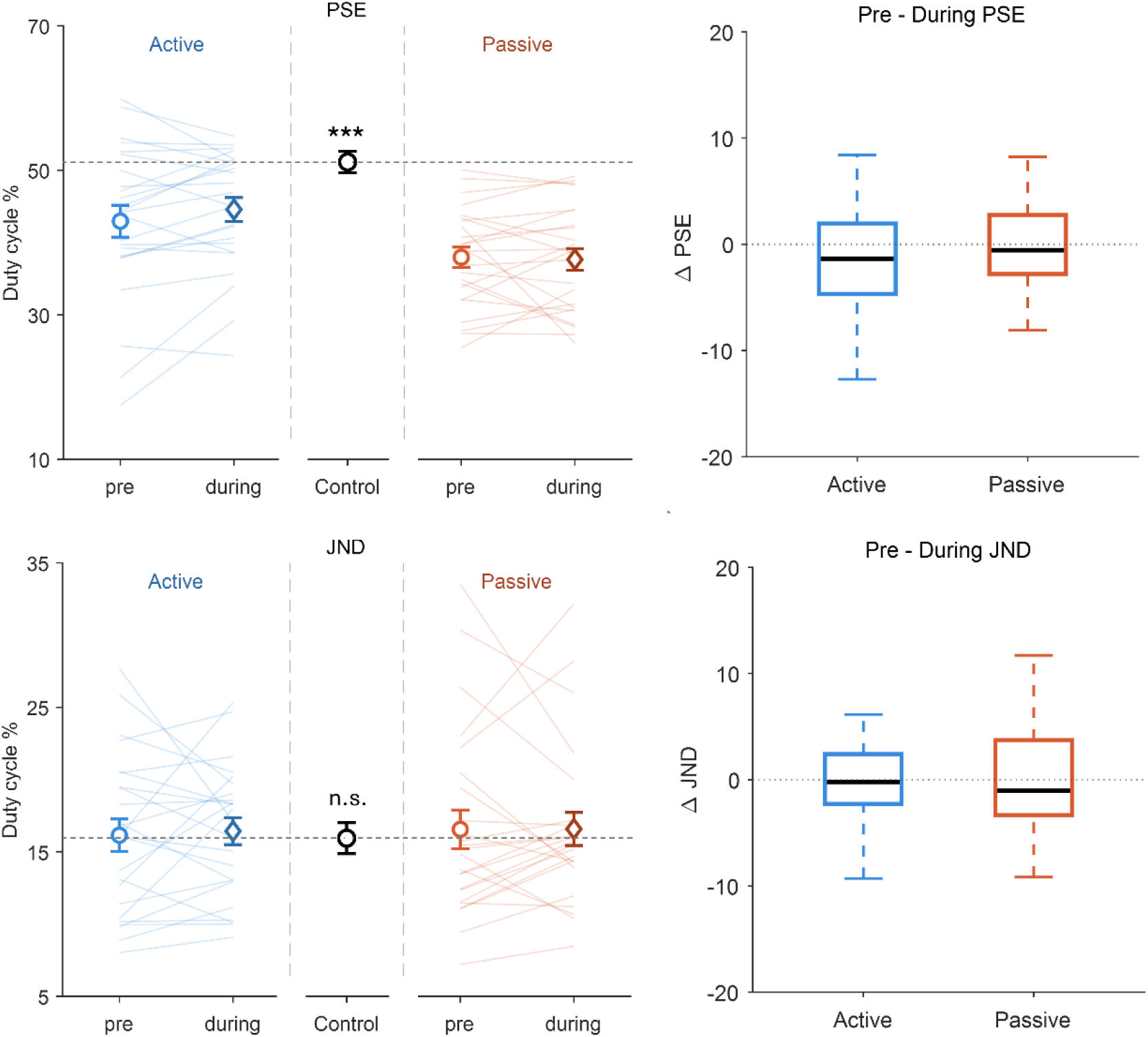
**Results**: Experiment 2 (pre-movement). Left panels show individual and mean PSE (top) and JND (bottom) values in duty cycle (%) for Active (blue) and Passive (orange) conditions across Pre-movement, During-movement, and Control. Thin lines connect individual participants. Error bars represent ±1 SEM. The dashed grey horizontal line indicates the reference duty cycle. Asterisks indicate significant deviation from Control (***p < .001); n.s. = not significant. Right panels show the Pre − During difference scores (Δ = Pre − During) for PSE (top) and JND (bottom). Boxplots show median, interquartile range, and whiskers to 1.5×IQR. Neither PSE nor JND delta differed significantly from zero in either movement context.

##### Active movement

A significant main effect was observed, F (2, 46) = 14.89, p < .001. Active Pre-movement (M = 4.30, SD = 1.08) yielded significantly lower PSEs than Control (M = 5.12, SD = 0.73), t (23) = 4.15, p = .001, d = 0.85. Active During-movement (M = 4.46, SD = 0.82) was also significantly lower than Control, t(23) = 4.09, p = .001, d = 0.84. PSEs did not differ significantly between pre-movement and during-movement, t (23) = −1.53, p = .139, d = −0.31. (Figure 4A)

##### Passive movement

A significant main effect was also observed, F (2, 46) = 46.20, p < .001. Both Passive Pre-movement (M = 3.80, SD = 0.69) and Passive During-movement (M = 3.77, SD = 0.74) differed significantly from Control (both p < .001). Crucially, PSEs did not differ between Passive Pre-movement and Passive During-movement (t(23) = 0.28, p = .783), indicating that the bias had reached its full extent before the arm was physically displaced. (Figure 4A)

To compare the temporal dynamics of bias across movement types, we computed for each participant the change in PSE from pre-movement to during-movement (Δ = Pre − During) separately for active and passive conditions. Neither delta differed significantly from zero (Active: t (23) = −1.53, p = .139, d = −0.31; Passive: t (23) = 0.28, p = .783, d = 0.06), and the two deltas did not differ from each other, t (23) = −1.09, p = .287, d = −0.22. Thus, although passive movement produced stronger overall bias, the magnitude of the pre-to-during change was comparable across movement types. (Figure 4B)

#### Precision (JND)

The 2 × 2 ANOVA on JND values revealed no significant effects: Condition, F (1, 23) = 0.09, p = .773; Time, F (1, 23) = 0.05, p = .823; interaction, F (1, 23) = 0.04, p = .839. The five-level one-way ANOVA similarly showed no effect, F (4, 92) = 0.13, p = .972. Mean JNDs were comparable across all conditions (range: 1.59 - 1.66), and no post-hoc comparison approached significance (all p = 1.000). Discrimination precision was unaffected by movement condition, movement phase, or their combination (Figure 4C).

Delta analyses confirmed no pre-to-during change in JND for either condition (Active: t(23) = −0.32, p = .756; Passive: t (23) = −0.03, p = .976), and the two deltas did not differ (t(23) = −0.21, p = .839). (Figure 4D)

#### Violating the predicted sensory consequences of the upcoming movement

The central claim of the present study is that the perceptual bias observed is driven by movement execution, rather than movement planning. If this interpretation is correct, then the bias should not be present when participants plan a movement but do not execute it, as in the NoGo condition. We tested this directly by comparing control-normalized PSE difference scores (Δ = Control − Condition) from the pre-movement conditions of Experiment 2 with the NoGo conditions of Experiment 1 (Figure 5A–B).

**Figure 5.**
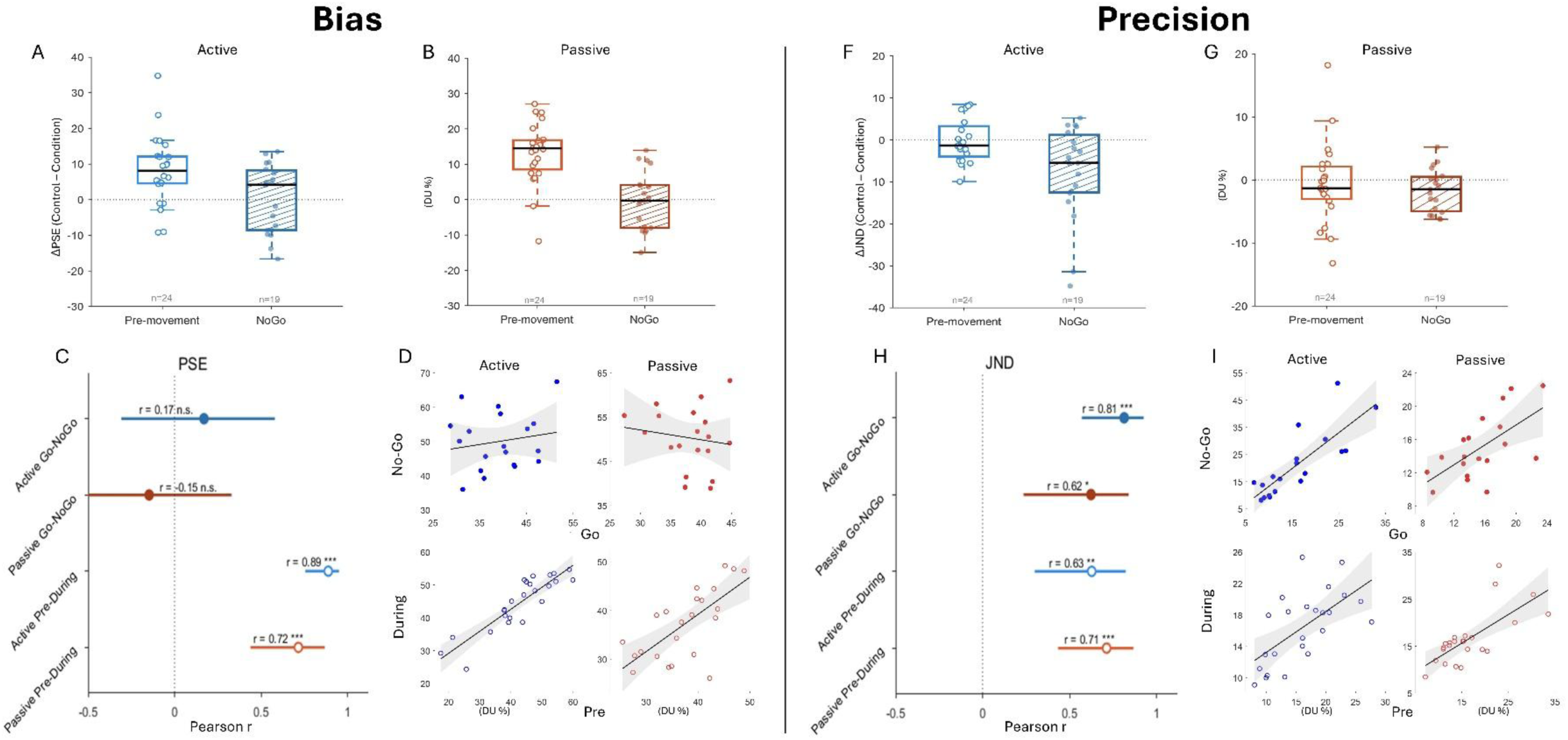
Cross-experiment comparison and rank-order stability of individual differences: A–B, F–G: Control-normalized difference scores comparing pre-movement trials from Experiment 2 with NoGo trials from Experiment 1. Difference scores were computed as Δ = Control − Condition. Panels A–B show ΔPSE for Active and Passive contexts; panels F–G show ΔJND for Active and Passive contexts. Open boxes indicate pre-movement conditions (n = 24), and hatched boxes indicate NoGo conditions (n = 19). Dots represent individual participants. The dashed horizontal line indicates zero, corresponding to no difference from baseline. C and H: Rank-order stability of individual differences across the theoretically critical paired conditions. Dots indicate Pearson correlation coefficients, and horizontal lines indicate 95% confidence intervals computed using Fisher’s z transformation. For PSE, rank-order stability was present across the Pre–During transition but absent across the Go–NoGo transition. For JND, rank-order stability was present across all critical comparisons. D and I: Scatter plots illustrating the corresponding individual-level correlations for PSE and JND. Filled markers show Go–NoGo correlations from Experiment 1; open markers show Pre–During correlations from Experiment 2. Blue indicates Active conditions and orange indicates Passive conditions. Regression lines are shown with 95% confidence bands. Asterisks indicate significant one-sample tests against zero, between-condition comparisons, or Holm-corrected correlation tests, as appropriate. *p < .05, **p < .01, ***p < .001. Full correlation matrices and additional pairwise comparisons are reported in the Supplementary Materials.

For PSE, one-sample t-tests showed that pre-movement deltas differed significantly from zero in both the active context (M = 0.820, SD = 0.967, t (23) = 4.152, p < .001) and the passive context (M = 1.320, SD = 0.860, t (23) = 7.521, p < .001). In contrast, NoGo deltas did not differ from zero in either context (Active NoGo: t (18) = −0.324, p = .749; Passive NoGo: t (18) = −0.506, p = .619). Direct between-experiment comparisons confirmed that pre-movement deltas were larger than NoGo deltas in both the active context (mean difference = 0.813, p = .009) and the passive context (mean difference = 1.342, p < .001). Thus, perceived intensity was reduced before movement onset only when movement was imminent, but not when movement was planned yet withheld.

For JND, the cross-experiment pattern was different (Figure 5F–G). Active NoGo produced significantly worse precision than Active Pre-movement (t (41) = 2.92, p = .006, d = 0.90), whereas Passive NoGo and Passive Pre-movement did not differ (t (41) = 0.65, p = .517). This indicates that cancelling an active motor plan selectively disrupted tactile precision, whereas withholding a passive displacement did not produce the same effect.

Rank-order stability of individual differences further supported this dissociation (Figure 5C–D, H–I). For PSE, strong stability was observed across the pre-to-during movement transition (Active Pre–During: r = .89, p < .001; Passive Pre–During: r = .72, p < .001), but not across the Go-to-NoGo transition (Active Go–NoGo: r = .17, n.s.; Passive Go–NoGo: r = −.15, n.s.). Thus, individual differences in bias were preserved across Pre–During conditions, but not across Go–NoGo conditions.

For JND, individual differences were stable across all critical comparisons. Significant correlations were observed for Active Go–NoGo (r = .81, p < .001), Passive Go–NoGo (r = .62, p < .05), Active Pre–During (r = .63, p < .01), and Passive Pre–During (r = .71, p < .001). Thus, PSE showed state-dependent rank-order stability, whereas JND showed stable individual differences across all critical motor-state comparisons.

## Discussion

In the present study, we demonstrated that tactile precision is predictively preserved during limb movement. Using a go/no-go paradigm, participants were required to withhold a planned movement unpredictably. Under these conditions, sensitivity to tactile stimulation intensity decreased markedly, even though the stimulated arm remained stationary. This finding indicates that tactile sensitivity is actively modulated during movement planning. Determining an object’s texture through manual exploration requires comparing incoming tactile stimulation with the speed of one’s own movement, which strongly influences receptor responses(24). Previous studies suggested that movement speed can be inferred directly from sensory input, as in passive stimulation conditions (25) (2). However, our findings suggest that, during active movement, this process is additionally supported by an efference copy.

On Active NoGo trials, participants planned a voluntary movement and generated a motor prediction; when the stop signal cancelled the movement, the predicted sensory consequences never arrived. This mismatch introduces noise into perceptual evaluation, degrading discrimination. On Passive NoGo trials, no efference copy is generated, no prediction error arises, and precision is unaffected. Equivalence testing (TOST) confirmed Passive Go and Passive NoGo JNDs were statistically equivalent. This interpretation is consistent with evidence that central predictive mechanisms modulate tactile processing independently of peripheral consequences (26) (27) (28). Considered alongside our previous work, this represents one of several ways the prediction–feedback loop can be disrupted (2, 3). The Active NoGo effect should nonetheless be interpreted with some caution, as cancelling a planned movement engages inhibitory control processes (29) (30) that can have their own somatosensory consequences(31) contributing to degraded discrimination. However, also in that case, movement planning, i.e. the cancellation of a movement, would influence tactile precision.

Experiment 1 showed that NoGo trials abolished the bias entirely. Despite full foreknowledge, motor planning, and strong expectation to move, perceived intensity remained equivalent to the stationary baseline. Experiment 2 further showed that, when movement occurred, the bias was already present before overt motion onset. Together, these findings indicate that pre-movement somatosensory gating reflects the influence of movement execution rather than movement planning, with pre- and during-movement bias reflecting the same execution-linked process at different time points.

This interpretation is consistent with classical accounts of reafferent masking (4, 7), recent evidence that tactile suppression is tightly coupled to movement onset (32), and findings showing somatosensory gating before movement when execution is inevitable (23). Unlike the expectancy effects reported by Voss et al. (2008) (16), movement planning was not sufficient to produce the reduction in perceived intensity. The same logic applies to passive movement: the bias emerged before passive displacement and disappeared when displacement was withheld. Because passive displacement generates no efference copy, the effect is most parsimoniously explained by movement-related sensory reafference (33).

### Individual differences reinforce the dissociation

The dissociation is supported not only at the group level but also by how the two measures behave at the individual level. Precision showed strong rank-order stability across every pair of related conditions in both experiments: participants who were imprecise in one state were imprecise in the others, regardless of motor execution, probe timing, or active-versus-passive context. Precision behaves as a stable, trait-like property of each participant consistent with a predictive mechanism that each individual carries across states of the motor system.

Across the pre-to-during-movement transition of Experiment 2, individual differences in bias were preserved, and they were preserved even across the active-versus-passive distinction every condition in that experiment shares one feature: movement actually occurred or was about to occur. Across the Go-to-NoGo transition of Experiment 1, bias showed no individual-level stability whatsoever. This asymmetry is the key observation: individual differences in bias carry over across pairs of conditions in which movement occurs in both states and vanish across pairs in which one state involves movement and the other does not. The absence of stability across Go and NoGo is therefore not a measurement-reliability failure, instead it is direct individual-level evidence that the bias mechanism is on-or-off with respect to movement execution, rather than planning.

### Two dissociable mechanisms

This dissociation aligns with the distinction drawn by Press et al. (2023) (1) between quasi-predictive gating mechanisms, which attenuate sensory processing on a moving effector regardless of specific predictions, and truly predictive mechanisms that track the match between predicted and actual sensory outcomes. In their framework, gating reflects reduced afference associated with efference and may operate at the spinal level, producing a non-specific attenuation tied to the occurrence of movement rather than to any particular prediction. Our PSE results fit this description: the bias is movement-locked, non-specific to active versus passive displacement, and absent when movement does not occur. Our JND results, by contrast, depend on whether a specific motor prediction was generated and fulfilled the hallmark of the truly predictive mechanism Press et al. distinguish from gating.

In conclusion, somatosensory gating is not a unitary phenomenon. The motor act itself produces the reduction in perceived intensity, consistent with a quasi-predictive gating mechanism tied to the occurrence of movement. The integrity of the motor prediction determines discrimination precision, consistent with a truly predictive mechanism that tracks the match between expected and actual sensory consequences. The motor system shapes tactile perception through at least two functionally distinct mechanisms: one tied to the physical consequences of movement and the other to the predictive processes that accompany it.

## Funding

none

## Competing Interests

The authors declare no competing interest.

## Ethics approval

This study was approved by the local ethics committee of the mathematical and natural science faculty at Heinrich-Heine-University Düsseldorf (Ethics-approval associated with ERC grant 757184).

## Consent to participate

All participants provided written informed consent prior to participation.

## Consent for publication

Not applicable.

## Availability of data and materials

The data is available in: https://doi.org/10.5281/zenodo.20647540

## Code availability

The code for the main analysis is available here: https://github.com/pierodonpac/Sensory-Gating-Analysis

## Author Contributions

All authors contributed to the study concept and to the design. Stimuli were designed by P.D.P and performed the data analysis. All authors contributed to the interpretation of results. P.D.P. drafted the manuscript, and E.Z. provided critical revisions. All authors approved the final version of the manuscript for submission.

## Notes

https://doi.org/10.5281/zenodo.20647540

### Competing Interest Statement

The authors have declared no competing interest.

